# Performance and efficiency of machine learning algorithms for analyzing rectangular biomedical data

**DOI:** 10.1101/2020.09.13.295592

**Authors:** Fei Deng, Jibing Huang, Xiaoling Yuan, Chao Cheng, Lanjing Zhang

**Author notes:** Correspondence: Lanjing Zhang, MD, Department of Pathology, Princeton Medical Center, 1 Plainsboro Rd, Plainsboro, NJ 08536, USA. or.

## Abstract

Most of the biomedical datasets, including those of ‘omics, population studies and surveys, are rectangular in shape and have few missing data. Recently, their sample sizes have grown significantly. Rigorous analyses on these large datasets demand considerably more efficient and more accurate algorithms. Machine learning (ML) algorithms have been used to classify outcomes in biomedical datasets, including random forests (RF), decision tree (DT), artificial neural networks (ANN) and support vector machine (SVM). However, their performance and efficiency in classifying multi-category outcomes in rectangular data are poorly understood. Therefore, we aimed to compare these metrics among the 4 ML algorithms. As an example, we created a large rectangular dataset using the female breast cancers in the Surveillance, Epidemiology, and End Results-18 (SEER-18) database which were diagnosed in 2004 and followed up until December 2016. The outcome was the 6-category cause of death, namely alive, non-breast cancer, breast cancer, cardiovascular disease, infection and other cause. We included 58 dichotomized features from ~53,000 patients. All analyses were performed using MatLab (version 2018a) and the 10-fold cross validation approach. The accuracy in classifying 6-category cause of death with DT, RF, ANN and SVM was 72.68%, 72.66%, 70.01% and 71.85%, respectively. Based on the information entropy and information gain of feature values, we optimized dimension reduction (i.e. reduce the number of features in models). We found 22 or more features were required to maintain the similar accuracy, while the running time decreased from 440s for 58 features to 90s for 22 features in RF, from 70s to 40s in ANN and from 440s to 80s in SVM. In summary, we here show that RF, DT, ANN and SVM had similar accuracy for classifying multi-category outcomes in this large rectangular dataset. Dimension reduction based on information gain will significantly increase model’s efficiency while maintaining classification accuracy.

## I. INTRODUCTION

Most of the biomedical data are rectangular in shape, including those of ‘omics, large cohorts, population studies and surveys. Few missing data were present in these datasets. An increasingly number of human genomic and survey data have been produced in recent years [1]. Rigorous analyses on these large datasets demand considerably more efficient and more accurate algorithms, which are poorly understood.

Machine learning (ML) algorithms are aimed to produce a model that can be used to perform classification, prediction, estimation or any other similar task [2, 3]. The unknown dependencies/associations are estimated based on a given dataset and later can be used to predict the output of a new system or dataset [2-4]. Therefore, ML algorithms have been used to analyze large biomedical datasets [5–7]. Studies have compared the accuracy of several ML algorithms for classifying microarray or genomic data [8–10], and show a superior performance of random forests (RF). However, only few studies accessed the accuracy of ML algorithms in classifying multi-category outcomes which are important for more-in-depth understanding of biological and clinical processes. Hence, we aimed to understand the performance and efficiency of RF, decision tree (DT), artificial neural networks (ANN) and support vector machine (SVM) algorithms in classifying multi-category outcomes of rectangular-shaped biomedical datasets.

As an example, we used a large population-based breast cancer dataset with long-term follow-up data to create a large rectangular dataset. The reasons were: 1. Cancer is one of most common diseases [11]. Knowledge gained by studying cancer can be easily generalized to other biomedical fields; 2. Breast cancer is the second most common cancer in the U.S. women [11, 12], and would provide a sufficiently large sample size (n>50,000); 3. Breast cancers diagnosed in 2004 had 12 years of follow up on average, and very few missing outcomes/follow-ups were anticipated; 4. Breast cancers diagnosed in 2004 would have a moderately-good prognosis and their outcomes will be rather diversified (i.e. many patients might not die of breast cancer). Therefore, this study is designed to systematically compare the performance and efficiency of DT, RF, SVM and ANN algorithms in classifying multicategory causes of death (COD) in a large biomedical dataset (breast cancers).

In this study,

## II. Methods

### 2.1 Data Analysis

We obtained individual-level data from the Surveillance, Epidemiology, and End Results-18 (SEER-18) (www.seer.cancer.gov) SEER*Stat Database with Treatment Data using SEER*Stat software (Surveillance Research Program, National Cancer Institute SEER*Stat software (seer.cancer.gov/seerstat) version <8.3.6>) as we did before [13–15]. SEER-18 is the largest SEER database including cases from 18 states and covering near 30% of the U.S. population. The SEER data were de-identified and publicly available. Therefore, this was an exempt study (category 4) and did not require an Institutional Review Board (IRB) review. All incidental invasive breast cancers of SEER-18 diagnosed in 2004 were included and had the follow-up to December 2016. The individual deaths were verified via certified death certificates (2019 data-release). We chose the diagnosis year of 2004 with the consideration of implementation of the 6th edition of the Tumor, Node and Metastasis staging manual (TNM6) of the American Joint Commission on Cancer (AJCC) in 2004. Moreover, we only included the primary-cancer cases that had a survival time > 1 month, age of 20+ years, and known COD.

The features (i.e. variables) were dichotomized for more efficiency and slightly better performance [7]. A total of 58 features were included (**Supplementary Table 1**). The outcomes of the classification models were the patient’s 6-category COD. The COD were originally classified using SEER’s recodes of the causes of death, which were collected through death certificates of deceased patients (https://seer.cancer.gov/). We simplified the SEER COD into 6 categories based on the prevalence of COD [16–18], including Alive, NonBreast cancer, Breast cancer, CVD, Infection and Other cause.

The most common task of ML techniques in learning process is classification [19]. The ten-fold cross-validation approach was used to tune all models, which is also termed as model optimization (See supplementary methods). We conducted experiments on a computer configured as follows: Huawei KLV-WX9 laptop; Windows 10 64-bit (DirectX 12); Intel Core i7-8565U @ 1.80GHz (Quad core); Motherboard of KLV-WX9-PCB(I/O – 9D84 for mobile 8th Gen Intel Core processor family); 8 GB Memory (SK Hynix LPDDR3 2133MHz); Hard disk drive of WDC PC SN720 SDAPNTW-512G-1127(512 GB / SSD); Graphics card of Nvidia GeForce MX250 (2 GB). All ML analyses were carried out using MATLAB (version 2018a, MathWorks, Natick, MA).

### 2.2 Model tuning

The detailed model tuning process is described in supplementary material. Several DT methods, such as CHAID, CART and exclusive CHAID are available with MATLAB [20]. We used CART (Classification and Regression Tree) to predict the categories, using the Gini index as split criterion and 100 iterations for each run. There are no default RF packages/toolboxs in MATLAB’s own toolbox. We thus used the Randomforest-matlab open source toolbox developed by Jaiantilal et al. [21, 22].

In tuning RF models, the parameter nTrees, which was to set how many trees in a random forest, and may have an impact on the classification results. We set the value of nTrees from 1 to 600 separately, the results show that if this parameter is not too few (greater than 10), the accuracy of recognition can reach 71% to 72% (**Supplementary Table 2**). Therefore, we set this parameter to 500 in RF-based analyses. The value of Mtry node for the best model performance was identified by setting the parameter value from 1 to 30 with 1 as the interval. We found 7 was the value, which was indeed consistent with the default value generated using the MATLAB model’s default (i.e. Mtry=floor(sqrt(size(P_train,2)))).

Among different training algorithms of ANN, we used the Trainscg algorithm because it is the only conjugate gradient method that did not require linear search. The number of input layer nodes is 58, and the number of output layer nodes is 6. To tune the ANN model, we conducted experiments of either single hidden or double hidden layers, with the node numbers ranged from 5 to 100. According to the tuning results of accuracy and mean squared error (MSE), we would set the model to double hidden layers, and the number of layers for the highest accuracy and lowest MSE.

We used the multi-class error-correcting output codes (ECOC) model the SVM modelling, which allows classification in more than two classes; and the MATLAB fitcecoc function that creates and adjusts the template for SVM [23]. The Kernel functions considered in the SVM were: Linear, Radial basis function, Gaussian and Polynomial.

### 2.3 Performance analysis

We analyzed the performance metrics of each proposed model, including accuracy, recall, precision, F1 score and specificity [24, 25]. They were defined as follows:

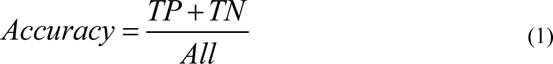

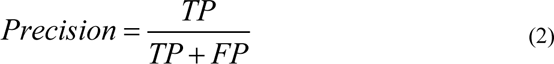

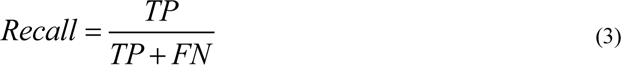

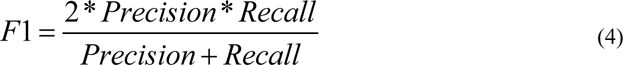

True Positive (TP) and True Negative (TN) were defined as the number of samples that are classified correctly. False Positive (FP) and False Negative (FN) were defined as the number of samples that are misclassified into the other mutational classes [24, 25]. The specificity or true negative rate (TNR) is defined as the percentage of mutations that are correctly identified:

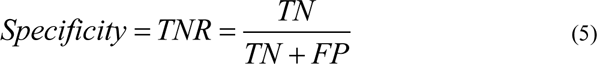

The receiver operating curve (ROC) is a graph where recall is plotted as a function of 1-specificity. It can more objectively measure the performance of the model itself [23]. The model performance was also evaluated using the area under the ROC, which is denoted the area under curve (AUC). An AUC value close to 1 highlight a high-performance model, while an AUC value close to 0 demonstrate a low model performance [26, 27]. AUC is independent of class prior distribution, class misclassification cost and classification threshold, which can more stably reflect the model’s ability to sort samples and characterize the overall performance of the classification algorithm. The formula used to determine the AUC can be written as follows (Hong et al., 2018a) [27, 28].

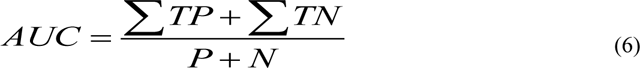

Where P is the total number of positive class and N is the total number of negative class.

### 2.4 Dimension reduction based on the information entropy and information gain

Information entropy is an indicator to measure the purity of the sample set. The formula is as follows:

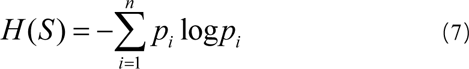

The measure of information gain is to see how much information a feature can bring to the classification system. The more information it brings, the more important the feature. Information gain *IG*(A) is the compute of the difference in entropy from start to end the set S is split on an attribute A, the information gain is defined as follows [29][50]:

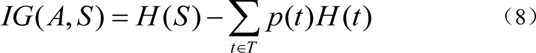

Where H(S) is the entropy of set S, T is the subsets created from splitting set S by attribute A such that *S* = *U*_*t∈T*_*t*, p(t) is the proportion of the number of elements in t to the number of elements inset S, and H(t) is the entropy of subset T.

The information gain of a feature can indicate how much information it brings to the classification system and can be used as a feature weight. When the model uses more features, the classification time will be longer. Arbitrarily reducing the characteristics will likely reduce classification accuracy. Therefore, we screened the features based on the calculated information gains to achieve a balance between run time and classification accuracy. We then step-wise deleted the features of the least information of gain, which were likely the least important. Because DT and RF were both ensemble-based algorithms and had similar performances, we only conducted dimension reduction with RF models and expect similar results with DT models.

## III. Results

### 3.1. Dataset characteristics and the model tuning

Of the 52,818 samples, there were missing values for tumor level (_grade) and level (lat_bi), of which there were 5,294 (~10%) vacancies in tumor grade (grade) and 352 (0.6%) vacancies in the level (lat_bi), and 317 (0.6%) data were missing at the same time. Here, we fill in with other invalid values, such as −5 to prefill these vacant feature values, and then act as input features for the model.

For DT models, the minimum number of leaf nodes with the best performance of the DT ranged 120-560 and bottomed at 359 (**Supplementary Fig. 1**). Therefore, the minimum number of samples contained in the leaf node was 359 for optimization. The overall classification accuracy could reach 72%, which was about 7% higher than the original DT, and the cross-validation error has also decreased. However, due to the uneven distribution of the data samples, some categories such as Non-Breast cancer, Infection and Other cause were pruned, causing loss of some data information.

For RF models, the parameter nTrees set how many trees in a RF and may have an impact on the classification results. The Mtry is the number of variables randomly sampled as candidates at each split and was optimized (**Supplementary methods**). After tuning the models, we set the nTrees parameter to 500 in RF-based analyses with the best Mtry node value of 7 (**Supplementary Table 2**).

According to the tuning results of accuracy and mean squared error (MSE) in ANN models, when the number of layers was greater than 20, the models’ performance appeared stabilized **(Table 1)**. Therefore, we set the model to double hidden layers, and the number of layers is 50.

For the SVM models of linear, radial basis function, Gaussian or polynomial function, we found the linear kernel function had the highest accuracy (71.87%) and shortest run-time (483.57s, **Table 2**), with a one-vs-one approach.

### 3.2 Performance analysis results

Based on the confusion matrices (**Fig. 1**), the 4 ML models appeared to have similar performance. The best classification accuracy of DT, RF, ANN and SVM models in this study were 72.68%, 72.66%, 70.01% and 71.85%, respectively, and seemed overall acceptable. However, to evaluate the pros and cons of a model, it is not enough to just look at the accuracy. The values of recall, TNR and F1 can specifically reflect the classification of each category. Further comparing the metrics among the four models show that the Recall and F1 in classifying Non-Breast cancer by RF model were significantly better than the DT, ANN and SVM models. In addition, Recall and F1 of Other cause were 1.83% and 3.38%, which were slightly higher than the DT and SVM models. This is mainly because that the RF model is an integrated learning algorithm; through the voting mechanism, one can balance a certain error. The accuracy of ANN model in classifying Alive, Non-Breast cancer, Breast cancer and CVD was not higher than those of the DT, RF model or SVM models. However, that of ANN model in classifying Other cause was the highest. Unfortunately, none of the above four models can effectively classify the COD of infection.

**Figure 1.**
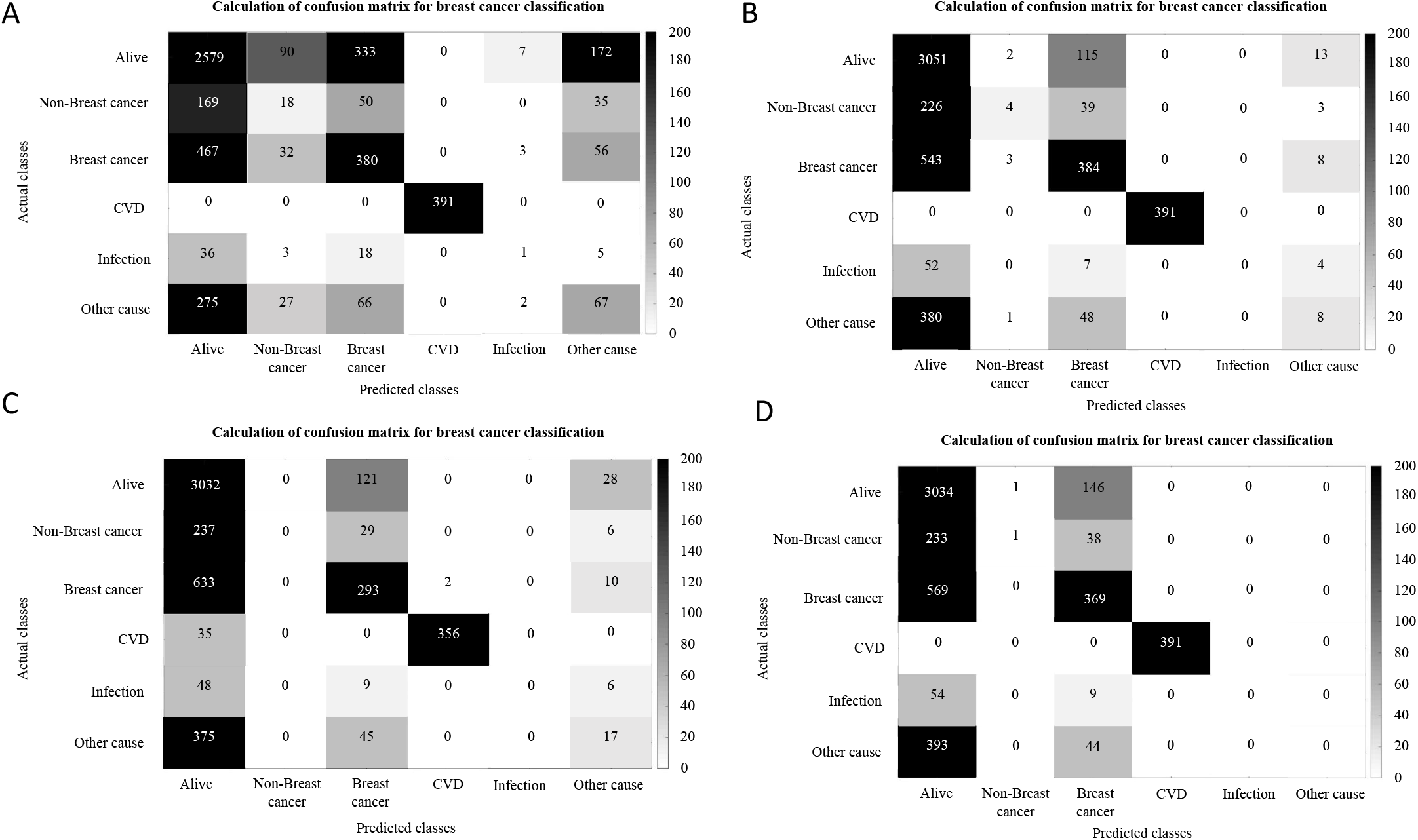
The confusion matrices of the tuned decision tree (A), random forest (B), artificial neural networks (C) and support vector machine (D) models.

### 3.3. ROC analysis results

After a comparative analysis of the above performance metrics, we found that the RF model was superior to the DT, ANN and SVM models. However, in the classification of Other causes, the ANN model has a higher recognition rate than the RF model, and the F1 values of Non-Breast cancer and Infection cannot be calculated. Therefore, we used ROC curve to further analyze Non-Breast cancer, Infection and Other cause in RF and ANN models.

The AUC of Non-Breast cancer, Infection and Other cause of the RF model are 0.6139, 0.7150 and 0.6217 respectively, which are better than 0.5481, 0.6373 and 0.6125 in the ANN model (**Fig. 2**). Because the AUC index can measure the performance of the model more objectively, the results show that the overall performance of the RF model is better than that of the ANN model.

**Figure 2.**
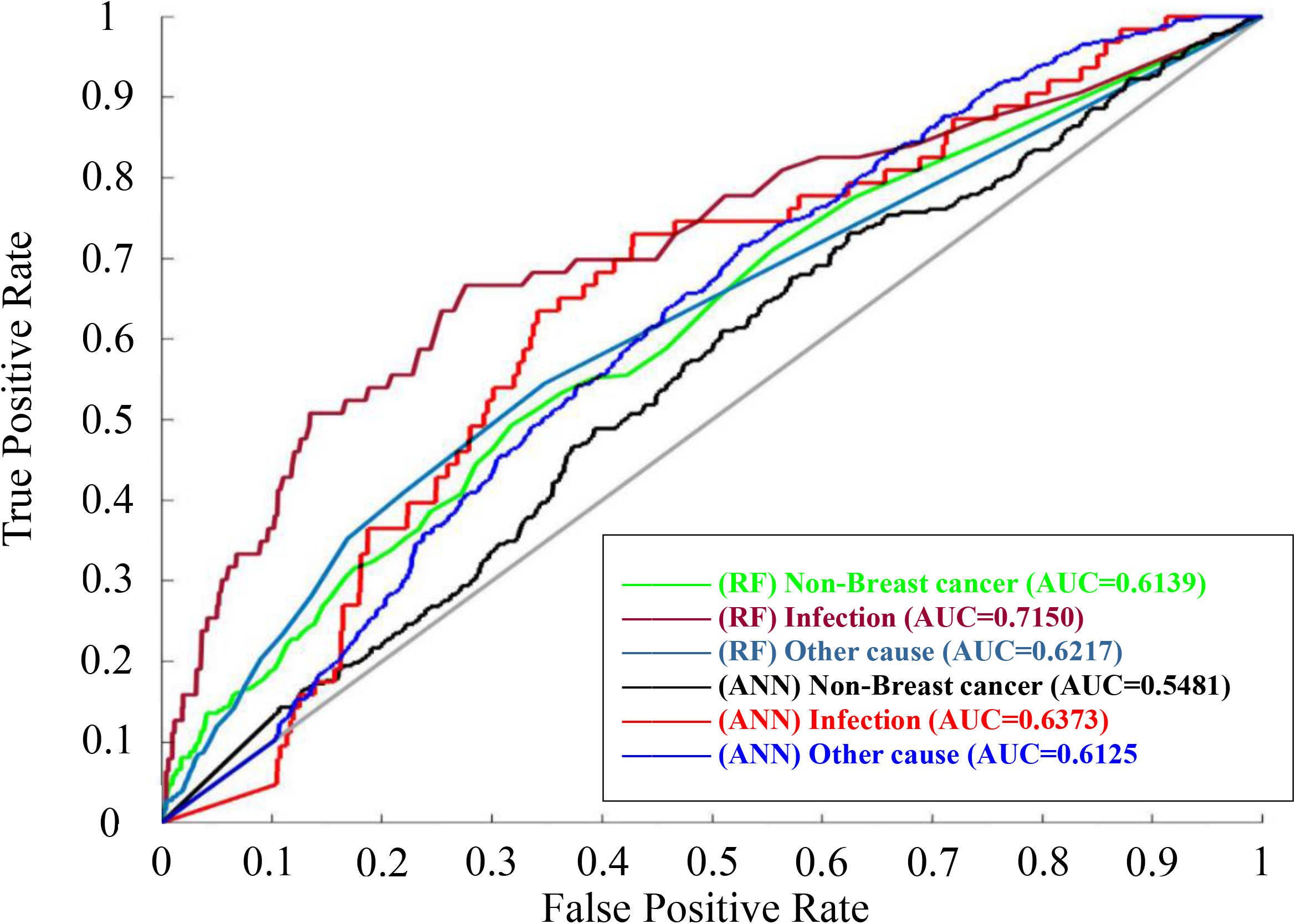
The receiver Operator Curve (ROC) of the tuned random forest and artificial neural networks models

### 3.4 Dimension reduction based on the information entropy and information gain

This dataset had 58 categorical/binary features, except the random id for grouping which was continuous. Based on the training datasets, the information entropy and information gain of 58 features were calculated **(Supplementary Table 3)**. Using information entropy and information gain, we obtained the following important features: random id for grouping; Cause of death-only cardiovascular; age >65; Surgery; TNM6 metastasis subgroup1; AJCC stage 4; TNM6 metastasis subgroup2; Surgery – other; TNM6 tumor subgroup1; TNM6 tumor subgroup2; AJCC stage 1; Surgery-lumpectomy; TNM6 lymph node subgroup5 and TNM6 lymph node subgroup1. Then we characterized the key features in a step-wise fashion (**Supplementary Fig. 2**).

We successfully reduced the data dimension based on information gain and shortened the run times in RF, ANN and SVM models, while maintaining the overall classification accuracy (**Table 4** and **Supplementary Tables 4-6**). Removal of feature with low information gain (0.0000-0.0005) in RF models led to slight increase in Alive and the overall accuracy rates, while no accuracy changes in CVD and Breast cancer classes. The classification of CVD was always 100%; Alive and the overall accuracy rate had a slight improvement, respectively about 0.8% and 0.3 %; Breast cancer and CVD had no significant changes; Non-Breast cancer accuracy was slightly reduced, and the classification effect is unstable, and the Infection category was still not recognized in the model due to the uneven distribution of data samples. Therefore, the features with low information gain (0.0000-0.0005) may be considered as redundant features, and deleted in the models, while the running times were scientifically reduced. We also found similar changes in ANN and SVM models.

## IV. Discussion

We here compared the performance and efficacy of DT, RF, ANN and SVM in classifying 6-category outcomes of a large rectangular database (58 features and ~53,000 samples). The accuracy in classifying 6-category COD with DT, RF, ANN and SVM was 72.68%, 72.66%, 70.01% and 71.85%, respectively. It is noteworthy that the accuracy in classifying 6-category outcomes is exponentially more difficult than classifying binary-category outcomes since it depends on 6 sequential classification processes (i.e. accuracy^6). Moreover, based on the information entropy and information gain of feature values, we could reduce the feature number in a model to 22 and maintained the similar accuracy, while the running time decreased from 440s for 58 features to 90s for 22 features in RF, from 70s to 40s in ANN and from 440s to 80s in SVM. The DT algorithm was not tested after dimension reduction for its lower performance than RF and its theoretical framework (ensemble-based) like RF.

Few studies to our knowledge investigated the dimension reduction of multicategory classification. Two-stage approach has been shown to effectively and efficiently select features for balanced datasets [30], but no specific reduction in run time was reported. We here show that dimension reduction and efficiency improvement can be achieved by removing features of low to medium information gain (<0.0005) in RF, ANN and SVM models, which apparently have little effect on the overall classification performance. Such a strategy may be applied to other ML models in classifying unbalanced large rectangular datasets, while caution should be used when classifying outcomes in a balanced dataset.

The four included ML algorithms each have their own theoretical frameworks. DT is a logical-based ML approach [31]. The structure of DT is similar to a flowchart. Using top-down recursion, the classification tree produces the category output. Starting from the root node of the tree, test and compare property values on its internal node, then determine the corresponding branch, and finally reach a conclusion in the leaf node of the DT. This process is repeated at each node of the tree by selecting the optimal splitting features until the cut-off value is reached [32]. A leafy tree tends to overtraining, and its test accuracy is often far less than its training accuracy. By contrast, a shallow tree can be more robust and be easy to interpret [33].

DT works by learning simple decision rules extracted from the data features. But RF are a combination of tree predictors such that each tree depends on the values of a random vector sampled independently, and with the same distribution for all trees in the forest [34]. When RF used for a classification algorithm, the deeper the tree is, the more complex the decision rules and the fitter the model [34, 35]. The generalization error of a forest of tree classifiers depends on the strength of the individual trees in the forest and the correlation between them. Random decision forests overcome the problem of over fitting of the DTs and are more robust with respect to noise.

ANN technique is one of the artificial intelligence tools. It is a mathematical model that imitates the behavior characteristics of animal neural networks [36]. This kind of network carries on distributed parallel information processing, by adjusting the connection between a large number of internal nodes, so as to achieve the purpose of processing information. After repeated learning and training, network parameters corresponding to the minimum error are determined, and the ANN model classifies the output automatically from the dataset.

SVM is another popular ML tool based on statistical learning theory, which was first proposed by Vladimir Vapnik and his colleagues [37, 38]. Unlike traditional learning methods, SVMs are approximate implementations of structural risk minimization methods. The input vector is mapped to a high-dimensional feature space through some kind of non-linear mapping which was selected in advance. An optimal classification hyperplane is constructed in this feature space, to maximize the separation boundary between the positive and negative examples [37, 38]. Support vectors are the data points closest to the decision plane, and they determine the location of the optimal classification hyperplane.

The four ML algorithms had different strengths and weaknesses. The RF algorithm in our study seems to have the best overall performance for its lack of being unable to classify some CODs (NA in the Table 3) and the best overall accuracy. Despite the similar classification accuracy (~72.66%), the DT algorithm could not accurately classify non-breast cancer group. Given the similar theoretical framework, we did not access its performance after dimension reduction. The ANN algorithm in our study is most efficient before and after dimension reduction. Surprisingly, we also notice a small increase in accuracy after dimension reduction which warrants further investigation. The SVM algorithm in our study appears very sensitive to the subgroup size (i.e. number of samples) and was not able to classify 2 of the 6 COD, although it also acceptable classification accuracy (71.8%).

The study’s limitations should be noted when applying our findings to other databases. First, this type of rectangular database is typical in survey and population-study, but not so in computational biology. The major difference is the large *p* in ‘omics datasets versus the large *n* in epidemiological datasets, which referred to feature number and sample number, respectively. Second, some of the outcomes were not accurately classified. It is likely owing to the unbalanced outcome distribution. On the other hand, such an undesired situation reflexes the real-world evidence/experience. Further studies are needed to improve the classification accuracy in the classes of fewer samples. Third, our works were exclusively based on MATLAB platform, and may not be applicable to other platforms such as R or python. Finally, ideally, we should use a large database to validate our models, but it is very difficult to curate and apply the tuned models to another large database that is similar to the SEER database.

In summary, we here show that RF, DT, ANN and SVM algorithms had similar accuracy for classifying multi-category outcomes in this large rectangular dataset. Dimension reduction based on information gain will significantly increase model’s efficiency while maintaining classification accuracy.

## Acknowledgement

We thank Lingling Han at Shenzhen Horb Technology Corporate, Ltd. for invaluable discussions and comments.

## Author contribution

FD, CC and LZ designed the study, FD and JH conducted the study and drafted the manuscript, all authors discussed, revised and edited the manuscript and LZ supervised the work.

