## Supplementary material for "Performance and efficiency of machine learning algorithms for analyzing rectangular biomedical data": Table 1

| **Table 1** Performance of artificial neural networks models with various numbers of layers | | | | |
| --- | --- | --- | --- | --- |
| Hidden layer settings | | Neural network algorithm | Total accuracy | MSE |
| Single hidden layer | 5 | trainscg | 60.29% | 0.0967 |
|  | 10 | trainscg | 60.28% | 0.1284 |
|  | 20 | trainscg | 70.19% | 0.0655 |
|  | 40 | trainscg | 70.22% | 0.0646 |
|  | 50 | trainscg | 70.21% | 0.0661 |
|  | 60 | trainscg | 70.22% | 0.0664 |
|  | 70 | trainscg | 70.17% | 0.0665 |
|  | 80 | trainscg | 60.28% | 0.1284 |
|  | 90 | trainscg | 70.02% | 0.0676 |
|  | 100 | trainscg | 70.21% | 0.0674 |
| Double hidden layers | [5,5] | trainscg | 63.87% | 0.0888 |
|  | [10,10] | trainscg | 68.65% | 0.0757 |
|  | [20,20] | trainscg | 70.20% | 0.0654 |
|  | [40,40] | trainscg | 70.18% | 0.0659 |
|  | [50,50] | trainscg | 70.01% | 0.0667 |
|  | [60,60] | trainscg | 63.93% | 0.0814 |
|  | [70,70] | trainscg | 70.14% | 0.0665 |
|  | [80,80] | trainscg | 60.28% | 0.1285 |
|  | [90,90] | trainscg | 70.22% | 0.0668 |
|  | [100,100] | trainscg | 68.48% | 0.0676 |

MSE: Mean squared error.
