## Supplementary material for "Performance and efficiency of machine learning algorithms for analyzing rectangular biomedical data": Table 2

| **Table 2** Classification accuracies and efficiencies of support vector machine models by kernel function | | | | | | | | |
| --- | --- | --- | --- | --- | --- | --- | --- | --- |
| Kernel function | Alive | Non-Breast cancer | Breast cancer | CVD | Infection | Other cause | Total  accuracy | Run time |
| Linear | 95.44% | 0.37% | 39.23% | 100.00% | 0.00% | 0.00% | 71.87% | 483.57s |
| Linear, Radial basis Function | 96.70% | 0.74% | 5.01% | 18.41% | 0.00% | 2.97% | 60.77% | 1562.80s |
| Gaussian | 96.70% | 0.74% | 5.01% | 18.41% | 0.00% | 2.97% | 60.77% | 697.61s |
| Polynomial | 19.49% | 7.72% | 42.43% | 100.00% | 15.87% | 40.27% | 30.59% | 6400.65s |
