## Supplementary material for "Performance and efficiency of machine learning algorithms for analyzing rectangular biomedical data": Table 3

**Table 3** The performance of decision tree, random forests, artificial neural networks and support vector machine models.

| Statistical  Measure | Alive | Non-Breast cancer | Breast cancer | | CVD | | Infection | Other cause |
| --- | --- | --- | --- | --- | --- | --- | --- | --- |
| **Decision tree** | | | | | | | | |
| Accuracy | 72.68% | | | | | | | |
| Precision | 71.83% | 0.00% | 62.52% | | 100.00% | | NA | 26.67% |
| Recall | 96.10% | 0.00% | 41.26% | | 100.00% | | 0.00% | 0.92% |
| Specificity (TNR) | 39.48% | 99.97% | 93.70% | | 100.00% | | 100.00% | 99.71% |
| F1 | 82.21% | NA | 49.71% | | 100.00% | | NA | 1.77% |
| **Random forests** | |  |  | |  | |  |  |
| Accuracy | 72.66% | | | | | | | |
| Precision | 71.75% | 40.00% | 64.76% | | 100.00% | | NA | 22.22% |
| Recall | 95.91% | 1.47% | 40.94% | | 100.00% | | 0.00% | 1.83% |
| Specificity (TNR) | 39.59% | 99.84% | 94.29% | | 100.00% | | 100.00% | 99.27% |
| F1 | 82.09% | 2.84% | 50.16% | | 100.00% | | NA | 3.38% |
| **Artificial Neural Networks** | | | | | | | | |
| Accuracy | 70.01% | | | | | | | |
| Precision | 69.54% | NA | | 58.95% | | 99.44% | NA | 25.37% |
| Recall | 95.32% | 0.00% | | 31.24% | | 91.05% | 0.00% | 3.89% |
| Specificity (TNR) | 33.40% | 100.00% | | 94.35% | | 99.94% | 100.00% | 98.73% |
| F1 | 80.41% | NA | | 40.84% | | 95.06% | NA | 6.75% |
| **Support vector machine** | | | | | | | | |
| Accuracy | 71.85% | | | | | | | |
| Precision | 70.84% | 50.00% | 60.89% | | 100.00% | | NA | NA |
| Recall | 95.39% | 0.37% | 39.34% | | 100.00% | | 0.00% | 0.00% |
| Specificity (TNR) | 37.86% | 99.97% | 93.53% | | 100.00% | | 100.00% | 100.00% |
| F1 | 81.30% | 0.73% | 47.80% | | 100.00% | | NA | NA |

NA, not applicable; TNR: true negative rate.
