## Supplementary material for "Performance and efficiency of machine learning algorithms for analyzing rectangular biomedical data": Table 4

| **Table 4. Information-gain based dimension reduction in the three machine learning models** | | | | | | | | |
| --- | --- | --- | --- | --- | --- | --- | --- | --- |
| Accuracy  Feature change | Alive | Non-Breast cancer | Breast cancer | CVD | Infection | Other cause | Total accuracy | Time, mean |
| **Random forests** |  |  |  |  |  |  |  |  |
| Raw data (58 features in model) | 95.92% | 1.47% | 40.94% | 100% | 0.00% | 1.83% | 72.66% | 440s |
| Delete IG<0.0001 (42 features in model) | 96.38% | 1.10% | 41.04% | 100% | 0.00% | 1.37% | 72.90% | 220s |
| Delete IG<0.005 (22 features in model) | 96.70% | 0.74% | 40.41% | 100% | 0.00% | 0.46% | 72.89% | 80s |
| Delete top three (18 features in model) | 95.82% | 0.00% | 41.26% | 0.00% | 0.00% | 0.00% | 65.03% | 60s |
| **Artificial neural networks** | | |  |  |  |  |  |  |
| Raw data (58 features in model) | 95.32% | 0.00% | 31.24% | 91.05% | 0.00% | 3.89% | 70.01% | 70s |
| Delete IG<0.0001 (42 features in model) | 95.43% | 0.72% | 43.65% | 99.72% | 0.00% | 0.23% | 72.62% | 70s |
| Delete IG<0.005 (22 features in model) | 95.75% | 0.00% | 42.62% | 100% | 0.00% | 1.14% | 72.71% | 45s |
| Delete top three (18 features in model) | 95.27% | 0.00% | 41.95% | 0.00% | 0.00% | 0.00% | 64.82% | 40s |
| **Support vector machine** | |  |  |  |  |  |  |  |
| Raw data (58 features in model) | 95.38% | 0.37% | 39.34% | 100% | 0.00% | 0.00% | 71.85% | 440s |
| Delete IG<0.0001 (42 features in model) | 95.47% | 0.37% | 38.38% | 100% | 0.00% | 0.92% | 71.81% | 120s |
| Delete IG<0.005(22 features in model) | 95.41% | 0.37% | 39.13% | 100% | 0.00% | 0.00% | 71.83% | 60s |
| Delete top three (18 features in model) | 95.82% | 0.37% | 39.02% | 0.00% | 0.00% | 0.00% | 64.65% | 80s |

IG, information gain.
